# Dietary glutamine supplementation alleviates age-related cardiac dysfunction by reducing elevated H3K27me3

**DOI:** 10.1101/2025.05.16.654624

**Authors:** Cassy Le, Pierre Coleman, Thomas Burgoyne, Jie Su, Helen R. Warren, Ravi Kumar, Rafael Oexner, Hyunchan Ahn, Angeliki Ntorla, Celio Santos, Ajay Shah, Mauro Giacca, Elisabeth Ehler, Joseph R. Burgoyne

**Affiliations:** School of Cardiovascular and Metabolic Medicine & Sciences, King’s College London; The British Heart Foundation Centre of Excellence, The Rayne Institute, St Thomas’ Hospital, London, SE1 7EH, UK; Electron Microscopy Unit, Royal Brompton Hospital, Sydney Street, London SW3 6NP, UK; National Heart and Lung Institute, Imperial College, Dovehouse Street, London SW3 6LR, UK; William Harvey Research Institute, Barts and The London School of Medicine and Dentistry, John Vane Science Centre, Queen Mary University of London, London, UK; NIHR Barts Cardiovascular Biomedical Research Centre, Barts and The London School of Medicine and Dentistry, Queen Mary University of London, London, UK; School of Cardiovascular and Metabolic Medicine & Sciences, King’s College London; The British Heart Foundation Centre of Excellence, James Black Centre, 125 Coldharbour Lane, London, SE5 9NU, UK; Department of Medicine I, University Hospital Würzburg, Oberdürrbacher- Straße 6, 97080 Würzburg, Germany; School of Cardiovascular and Metabolic Medicine & Sciences, King’s College London; The British Heart Foundation Centre of Excellence, Randall Centre for Cell and Molecular Biophysics, New Hunt’s House, London, SE1 1UL, UK

## Abstract

**Background:** Changes to the epigenetic landscape play an important role in cardiovascular aging, where alterations in histone modifications influence gene expression by regulating DNA accessibility and chromatin structure. Our investigation into epigenetic changes during myocardial aging revealed that the repressive epigenetic mark H3K27me3 is significantly upregulated in aged mice and humans. This increase in H3K27me3 was shown to impair cardiomyocyte autophagy and drive metabolic reprogramming, key features of myocardial aging and causatively linked to dysfunction. Notably, by alleviating this repressive mark through a modified diet, we successfully mitigated the aged myocardial phenotype.

**Methods:** Heart tissue from young and aged mice and humans was analyzed for H3K27me3 levels using immunoblotting and immunofluorescence staining. Genes regulated by H3K27me3 were identified through CUT&RUN-Seq and RNA-Seq, while metabolites were profiled using metabolomics. In neonatal rat ventricular myocytes (NRVMs), H3K27me3 levels were elevated by siRNA-mediated knockdown of UTX. Cellular metabolism was investigated using a Seahorse analyzer in cardiomyocytes with basal or elevated H3K27me3 levels. In further human studies, we assessed how circulating glutamine levels associate with the incidence of heart failure and the association of genetic variants within the *SLC1A5* region with heart disease. In aged mice, H3K27me3 levels were reduced through a modified diet, and heart function was evaluated using echocardiography. Subsequently, hearts were processed for biochemical analysis, and autophagy was assessed using electron microscopy.

**Results:** H3K27me3 was significantly elevated in the aged mouse and human myocardium. This observed elevation in H3K27me3 was found to be attributed to impaired glutamine metabolism, resulting from reduced expression of the glutamine transporter SLC1A5 in the aged myocardium. Furthermore, elevation in H3K27me3 was found to contribute to impaired cardiomyocyte autophagy and metabolic dysfunction. In aged mice supplemented with a high glutamine diet this attenuated myocardial H3K27me3 and improved cardiac function. Furthermore, a high-glutamine diet reversed H3K27me3-mediated impairment in cardiac autophagy in aged mice.

**Conclusions:** Reduction in SLC1A5 during aging is likely to lead to increased myocardial H3K27me3 that results in impaired autophagy and metabolic reprogramming that contribute to the aged cardiac phenotype. Our findings also suggest glutamine may improve cardiac health in the aged population by lowering H3K27me3.

**Clinical Perspective:** *What is new?:*

- Elevation in H3K27me3 during aging is likely to lead to impaired myocardial autophagy and metabolic reprogramming.
- Reduction in SLC1A5 during aging in humans is likely to contribute to myocardial dysfunction
- Dietary supplementation of glutamine improves cardiac function in aged mice.

*What are the clinical implications?:*

- Targeting H3K27me3 in the aged population may to mitigate cardiovascular disease
- Dietary glutamine supplementation offers a promising strategy to improve cardiac function during aging.

## Introduction

Over the past decade, increasing evidence has shown that progressive alterations in the epigenetic landscape are closely associated with aging. Functional studies have demonstrated that these epigenetic changes significantly influence the aging process^1-4^. It is now well established that lifespan is largely shaped by diet and other environmental factors that modulate epigenetic information. This aligns with findings from model organisms, where inhibitors of epigenetic enzymes profoundly impact health and lifespan^3,5-11^.

One of the most well-studied repressive marks is H3K27me3, a histone lysine modification primarily mediated by the polycomb repressive complex 2 (PRC2), which contains the catalytic subunits EZH1 or EZH2. To a lesser extent, H3K27me3 is also regulated by the histone-lysine N-methyltransferase EHMT2. The removal of the tri-methyl group from H3K27 is carried out by the H3K27-specific JmjC-domain demethylases UTX and JMJD3. The distribution of H3K27me3 plays a critical role in regulating autophagy, a key catabolic process essential for maintaining cardiovascular health^12^. Autophagy sustains cellular homeostasis by removing redundant and dysfunctional biomolecules and organelles, while also providing bioenergetic intermediates during hypoxia and nutrient deprivation^13^. Alterations in H3K27me3 impacts on autophagic flux, likely through the regulation of autophagy-related gene expression^14-18^.

During aging, the heart undergoes homeostatic imbalance, fibrosis, left ventricular wall thickening, and increased left atrial size^19^. These changes contribute to a heightened risk of cardiovascular diseases, including cardiac hypertrophy, atrial fibrillation, and heart failure. In this study, we observed a substantial elevation of H3K27me3 in the aged mouse and human myocardium. This modification was found to suppress autophagy and contribute to metabolic reprogramming associated with the aging cardiac phenotype^12^. Furthermore, limiting the increase in H3K27me3 through dietary glutamine supplementation preserved cardiac function and improved autophagy in aged mice.

## Methods

### Cell culture

Neonatal rat ventricular myocytes (NRVMs) were isolated from 1-3-day-old Sprague-Dawley rats (Charles River). Following cervical dislocation, hearts were excised, washed in ice-cold ADS buffer (116 mM NaCl, 20 mM HEPES pH 7.35, 1 mM NaH_2_PO_4_, 5.6 mM glucose, 5.4 mM KCl, 0.8 mM MgSO_4_), and ventricles were isolated and minced. Tissue digestion was performed using collagenase type 2 (0.8 mg/mL, Worthington) and pancreatin (0.6 mg/mL, Sigma) for 5-6 cycles (15 min each, 37°C). Cells were filtered, pelleted (1250 rpm, 5 min), and pre-plated on T175 flasks (1h, 37°C, 5% CO_2_) in plating medium (DMEM supplemented with 5% FBS, 10% horse serum, 1% non-essential amino acids, 1% penicillin/streptomycin/glutamine). NRVMs were then transferred to gelatin-coated plates at 5×10^5^ cells/well (6-well) or 2.5×10^5^ cells/well (12-well). After 24h, medium was replaced with maintenance medium (DMEM containing 20% Medium 199, 1% penicillin/streptomycin/glutamine, 0.1 mM BrdU) and refreshed every 48h.

### Glutamine deprivation

Cells were maintained in media with no glucose and no glutamine (DMEM, ThermoFisher, cat. A1443001), supplemented with 5% FBS and penicillin/streptomycin (ThermoFisher, cat. #15140122) for 24 hrs. L-glutamine (ThermoFisher, cat. #25030081) was then added to a final concentration of 0.1 mM, 0.5 mM, 2 mM and 4 mM and replenished daily for 4 days.

### Transfections

siRNA transfections were performed using Lipofectamine RNAiMAX (Invitrogen, cat. #13778030) as per manufacturer’s instructions. The following siRNAs were used at the following concentrations: Scrambled non-targeting siRNA (Dharmacon, D-001810-01-05, 100 nM), KDM6A (UTX) (Dharmacon, J-115517-08-0005, 100 nM) Slc1a5 #11 and #12 (Dharmacon, J-094416-11-0002 and J-094416-12-0002 respectively, 100 nM).

### Western blotting

Protein samples were extracted using sample buffer (50mM Tris-HCl pH 6.8, 2% SDS, 10% glycerol, 0.0025% bromophenol blue, 5% β-mercaptoethanol). Young and aged mouse cardiac tissues were obtained from The Jackson Laboratory (#000664) or from mice sacrificed after the in-vivo study. Human cardiac tissue from the left ventricle was provided by the Sydney Human Heart Bank at the University of Sydney. Human tissue was used in accordance with the ethical guidelines of King’s College London (REC reference LRS-23/24-39407), in compliance with current UK legislation. Protein samples were separated on 4-15% TGX or 4-12% Bis-Tris gels and transferred to PVDF membranes. Membranes were blocked with 10% non-fat milk in PBST, then incubated with primary antibodies overnight at 4°C. After washing, membranes were incubated with HRP-conjugated secondary antibodies for 1 hour at room temperature. Protein bands were visualised using chemiluminescent substrate. All samples were heated at 95°C for 5 minutes and stored at -20°C until analysis. Samples were immunoblotted using the following antibodies from Cell Signaling Technology: histone H3 (4499S), H3K27me3 (9733S), H3K4me3 (9751T), H3K9me3 (13969T), H3K36me3 (4909T), H3K79me3 (4260T), UTX (33510S), p62 (5114S), LC3 A/B (12741S), EZH1 (42088S), EZH2 (5246S), JMJD3 (3457S), EHMT2 (3306S) and SLC1A5 (8057S). were used as loading controls. HRP-conjugated secondary antibodies were used for detection (7074S).

### Immunofluorescence

Human cardiac tissue microarrays (Biomax Inc, HEN241a) and paraffin-embedded young (12-week) and aged (68-week) mouse heart sections (Jackson Laboratory, #TSH0022) were immunostained for H3K27me3. After deparaffinisation and rehydration, antigen retrieval was performed using Tris-EDTA buffer (pH 9.0) for human samples or sodium citrate buffer (pH 6.0) for mouse samples. Sections were blocked with 10% normal goat serum, incubated with primary antibody (H3K27me3, Cell Signaling, 9733S) overnight at 4°C, and visualised with Alexa Fluor-conjugated secondary antibody. Fluorescent imaging was performed using a NanoZoomer S60 scanner. H3K27me3 nuclear puncta were quantified using ImageJ for human samples and QuPath for mouse samples, with results expressed as percentage of total nuclear area.

### Circulating glutamine analysis

To investigate how circulating glutamine levels affect the incidence of heart failure (HF) in humans we analysed data from UK Biobank, a large-scale prospective population-based cohort. Glutamine levels were available for approximately 280,000 participants enrolled in the UK Biobank, and measured by Nightingale Health Ltd. using their high-throughput proton nuclear magnetic resonance (^1^H-NMR)-based metabolic biomarker profiling platform

We excluded individuals with a diagnosis of HF prior to recruitment or with glutamine values more than 5 standard deviations above or below the mean. Diagnosis of HF was defined as either (i) self-reported “HF/pulmonary oedema” or “cardiomyopathy”; or (ii) hospital inpatient or death registry records containing an International Classification of Diseases (ICD)-10 or ICD-9 code for heart/ventricular failure or cardiomyopathy (hypertrophic cardiomyopathy was not considered, definitions were adapted as previously defined elsewhere)^20,21^.

We fitted two Cox-proportional hazards (PH) models; one looking into glutamine level as a continuous variable and the other using a binary variable comparing participants with the lowest 10% of glutamine levels to the rest of the cohort. We adjusted both models for age and sex.

### GWAS study

We looked-up the results of the *SLC1A5* gene region (chr 19: 46,774,883 - 46,788,594; build 38) within a recent Heart Disease GWAS^22^, which included N=71,808 cases vs N=357,986 controls of European ancestry from the longitudinal cohort from the Department of Veterans Affairs Million Veteran Program, which used Electronic Health Record data to test the “other chronic ischemic heart disease” phenotype according to PheCode 411.8. A locus-zoom plot was created using the LocusZoom software^23^ covering the *SLC1A5* gene region ±500kb.

### Metabolomics

Lyophilised cardiac tissue samples from young and aged mice underwent metabolite extraction using a methanol/chloroform/water protocol. The initial extraction used methanol/chloroform (2:1 v/v) with sonication, followed by a secondary extraction with methanol/water. Combined supernatants were vacuum-concentrated, reconstituted in chloroform:methanol (1:3:3), and the polar metabolite fraction was isolated by centrifugation. LC-MS analysis was performed using a 1290 Infinity II UHPLC system coupled with a 6546-Q-TOF mass spectrometer (Agilent Technologies).

### RNA-seq

Total RNA was extracted from young and aged mouse cardiac tissue using RNeasy mini kit with on-column DNase digestion (Qiagen). RNA quality was assessed by NanoDrop spectrophotometry. RNA-seq libraries were prepared using rRNA depletion and sequenced on an Illumina NovaSeq platform (2×150bp, 20M paired-end reads/sample). Sequencing reads were trimmed with Trimmomatic, aligned to the mouse genome (mm39) using STAR aligner, and unique gene hit counts calculated with featureCounts. Differential expression analysis was performed using DESeq2, with significance defined as adjusted p<0.05 and |log2FC|>1. Gene ontology enrichment was determined by Fisher’s exact test (p<0.05).

### CUT&RUN-seq

CUT&RUN was performed on young and aged mouse cardiac tissue using the Cell Signaling Technology CUT&RUN assay kit. Tissue was fixed with 0.1% formaldehyde, disaggregated, and processed according to manufacturer’s protocol. DNA was purified using spin columns from enriched chromatin samples. Libraries were prepared using NEBNext Ultra II DNA library prep kit and sequenced on Illumina NovaSeq 6000 (2×150bp paired-end). Raw reads underwent quality control with fastQC, adapter trimming with TrimGalore, and alignment to mouse genome (mm39) using bowtie2 with parameters for paired-end alignment. PCR duplicates were assessed using Picard tools. Peaks were called using MACS3 with a q-value threshold of 0.05. Genomic annotations were performed using ChIPseeker in R. Gene ontology analysis was created using ShinyGO 0.82.

### Animal study

All procedures were conducted in accordance with the Animals (Scientific Procedures) Act, 1986 (UK Home Office) and approved by King’s College London Animal Welfare Ethical Review Body. Male C57BL/6 mice were obtained from Charles Rivers at 12-weeks (young) and 64-weeks (aged), with aged mice maintained until 75-weeks before study initiation. Mice were housed in individually ventilated cages with controlled temperature and humidity, 12-hour light/dark cycle, and ad libitum access to food and water.

Three experimental diets were sourced from SAFE Nutrition Services (France). All mice were acclimated to the control diet for 3 weeks before randomisation into three groups: control diet, high amino acid (AA) diet, or high glutamine (Gln) diet. Both high AA and high Gln diets contained reduced pregelatinised cornstarch (152.6 g/kg vs. 352.6 g/kg in control). In the high Gln diet, the 200 g/kg cornstarch reduction was replaced with L-glutamine, while the high AA diet incorporated equivalent increases in various other amino acids, serving as an important control for nitrogen intake and reduced carbohydrate content.

Following the two-month intervention period and final echocardiography, mice were sacrificed, and hearts were harvested. After gentle removal of blood from the chambers, heart weight and tibia length were recorded. A small section from each apex was reserved for electron microscopy, with the remaining cardiac tissue flash-frozen in liquid nitrogen and stored at -80°C for subsequent analyses. No animals were excluded from this study.

### Echocardiography

Echocardiography was performed using high-frequency linear array transducers (MX400, 30MHz) with a Vevo 3100 imaging system (FUJIFILM VisualSonics). Mice were anesthetised with 2% isoflurane to maintain heart rates between 400-500 bpm and placed supine on a heating pad (37°C) with ECG monitoring. Measurements were taken before dietary intervention and at 4-week intervals following that.

Standard two-dimensional examinations were conducted to evaluate maximum LV dimensions in parasternal long-axis view. Images were archived as 300-frame cine loops in DICOM format for blinded offline analysis using Vevo Lab (v5.7.1).

### Electron microscopy

Heart tissue samples were fixed in Karnovsky EM fixative (2% PFA, 2% glutaldehyde in 0.1M cacodylate buffer, pH 7.4), washed, and stored in cacodylate buffer at 4°C. The samples were then incubated in 1% aqueous osmium tetroxide for 1 h at RT before en bloc staining in undiluted UA-Zero (Agar Scientific) for 30 min at room temperature (RT). Subsequently, the samples were dehydrated using an increasing ethanol concentration series (50, 70, 90, 100%), followed by a mixture of propylene oxide and Epon resin (1:1) for 1 h at RT. The cells were incubated in Epon 4h at RT, before embedding in fresh Epon for 24 hours at 60 °C. 100nm ultrathin sections were cut and images acquired on a JEOL 1400Plus transmission EM fitted with a Gatan Orius SC1000B charge-coupled device camera.

### Seahorse Assay

Mitochondrial function was assessed in NRVMs using the Seahorse XF Mito Stress Test (Agilent Technologies). Cells were seeded at 15,000/well in XF96 plates. The assay was performed in XF DMEM medium supplemented with glucose, pyruvate, and glutamine. Oligomycin, FCCP, and rotenone/antimycin A were sequentially injected to evaluate mitochondrial respiration parameters. To assess fuel source utilisation, separate wells received either vehicle, etomoxir, UK5099, or BPTES. Oxygen consumption rate was measured at baseline and following each injection. Data were normalised to cell number determined by Hoechst 33342 staining.

### SAM Assay

SAM levels were quantified in young and aged mouse cardiac tissue using the Bridge-It S-adenosyl methionine fluorescence assay kit (Mediomics, 1-1-1003) according to manufacturer’s protocol. The assay utilises the increased affinity of MetJ protein for its DNA-binding site in the presence of SAM, bringing fluorophore-conjugated components into proximity to generate a measurable fluorescence signal. SAM concentrations were normalised to total protein content determined by BCA protein assay (ThermoFisher, 23225).

### Statistical analysis

Data are shown as mean, with the standard error of the mean (SEM) represented by the error bars. The statistical tests include Student’s T-test, Mann-Whitney test, or ANOVA with multiple comparisons (Kruskal-Wallis or Turkey’s multiple comparisons) as specified in each figure legend. Results were considered significant at a 5% significance level.

## Results

### Elevated H3K27me3 in the aged mouse and human myocardium

The histone modifications H3K27me3, H3K4me3, and H3K9me3 were significantly elevated in the aged mouse myocardium (Fig. 1A). However, only elevation in H3K27me3 was sustained throughout aging (Fig. 1B), and was specifically observed in the heart and brain (Fig. S1A). In addition, increased H3K27me3 was found to be widespread across the aged mouse myocardium (Fig. 1C, Fig. S1B). These findings were corroborated in human tissue, where an age-dependent increase in histone H3K27me3 was detectable in homogenates and microarrays of human heart tissue (Fig. 1D, E). Collectively, these results identify elevated H3K27me3 as an epigenetic process that may contribute to myocardial aging.

**Figure 1.**
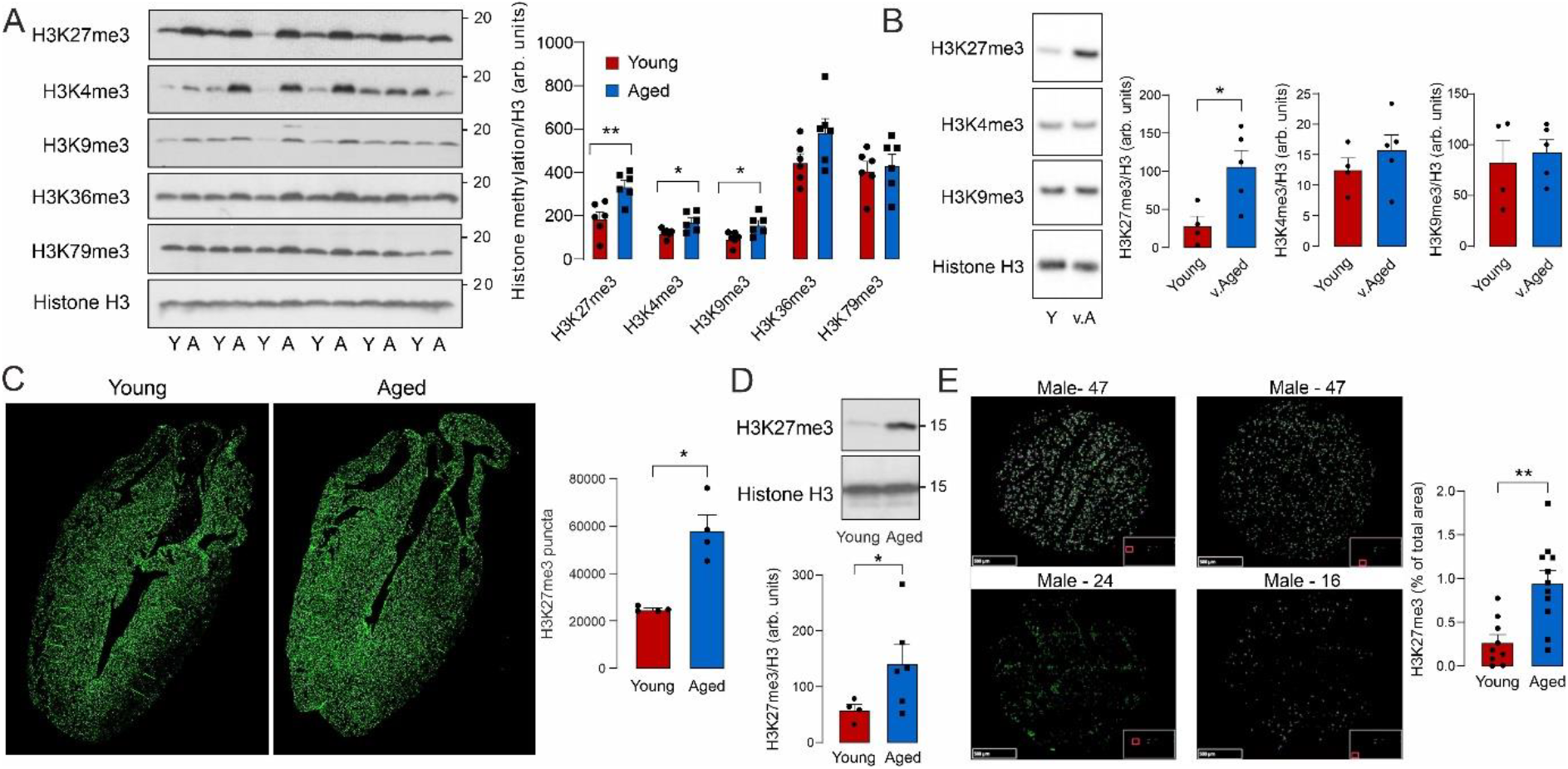
H3K27me3 is elevated in the aged mouse and human myocardium. A) Histone methylation in young (12-weeks) and aged mouse myocardium (68-weeks) (n=6). B) Histone H3K27me3, H3K4me3 and H3K9me3 in young (12-weeks) and very aged mice (85-weeks) (n=4-5). C) H3K27me3 visualised throughout the young and aged mouse heart (n=4). D) H3K27me3 in young (≤40 years) and aged human (>40 years) myocardium observed by immunoblotting of tissue samples (n=4-6) (E) or immunofluorescence staining of cardiac tissue microarrays (n=9-10) (F). *P<0.05; **P<0.01; ***P<0.005 by Student *t* test.

### Impaired glutamine metabolism due to loss in SLC1A5 is likely to underlie the aged-dependent increase in H3K27me3

The causes of the age-dependent increase in myocardial H3K27me3 were further investigated. This increase was found to be independent of the expression of methyltransferases (EZH1, EZH2, and EHMT2) or demethylases (JMJD3 and UTX) (Fig. 2A, B). Therefore, we next assessed the abundance of S-adenosyl methionine (SAM), the substrate required for histone methylation. However, the levels of this metabolite remained unchanged during aging (Fig. 2C). Since the H3K27me3 demethylases JMJD3 and UTX use alpha-ketoglutarate (α-KG) as a cofactor, we hypothesized that changes in α-KG levels might be responsible for the age-associated increase in H3K27me3. Metabolomic analysis revealed a loss of α-KG in the aged mouse myocardium (Fig. 2D). Additionally, glutamine (and glutamic acid), which can be converted to α-KG, was significantly reduced with aging (Fig. 2E, F). Based on these findings, the role of glutamine in maintaining myocardial health in humans was evaluated. Patients with lower circulating levels of glutamine exhibited a higher incidence of heart failure (hazard ratios per standard deviation of glutamine: 0.91 [0.89-0.93] and for the bottom decile: 1.37 [1.29–1.45]; both p < 2e-16) (Fig. 2G), further supporting its importance in preserving myocardial function. To further investigate the age-related decline in myocardial glutamine levels seen in mice, we examined the expression of the glutamine transporter SLC1A5. We found that SLC1A5 expression was significantly decreased in both the aged mouse and human myocardium (Fig. 2H-J). Supporting the role of this mechanism in glutamine depletion, knockdown of SLC1A5 in cardiomyocytes led to increased levels of histone H3K27me3 (Fig. 2K). Furthermore, the importance of *SLC1A5* in maintaining myocardial health was further supported by results from genome wide association studies (Fig 2L). A locus zoom plot revealed a clear signal of association with heart disease clustered around the *SLC1A5* gene^22^. The most significantly associated single nucleotide polymorphism near this gene is rs56010181 (at 46,695,675 bp position, build 38, on chromosome 19): the minor allele C (freq 12%) increases the odds of heart disease by OR=1.04 (p-value = 5.91×10^-8^). Within the *SLC1A5* gene interval itself, the most significantly associated SNP is rs559471437 (at 46,775,918 bp position, build 38, on chromosome 19): the minor allele T (freq 1.4%) increases the odds of heart disease by OR=1.12 (p-value = 6.02×10^-5^). In summary, these findings suggest that the loss of glutamine transport through SLC1A5 contributes to the increased H3K27me3 levels in the aged myocardium, as well as likely contributing to myocardial dysfunction.

**Figure 2.**
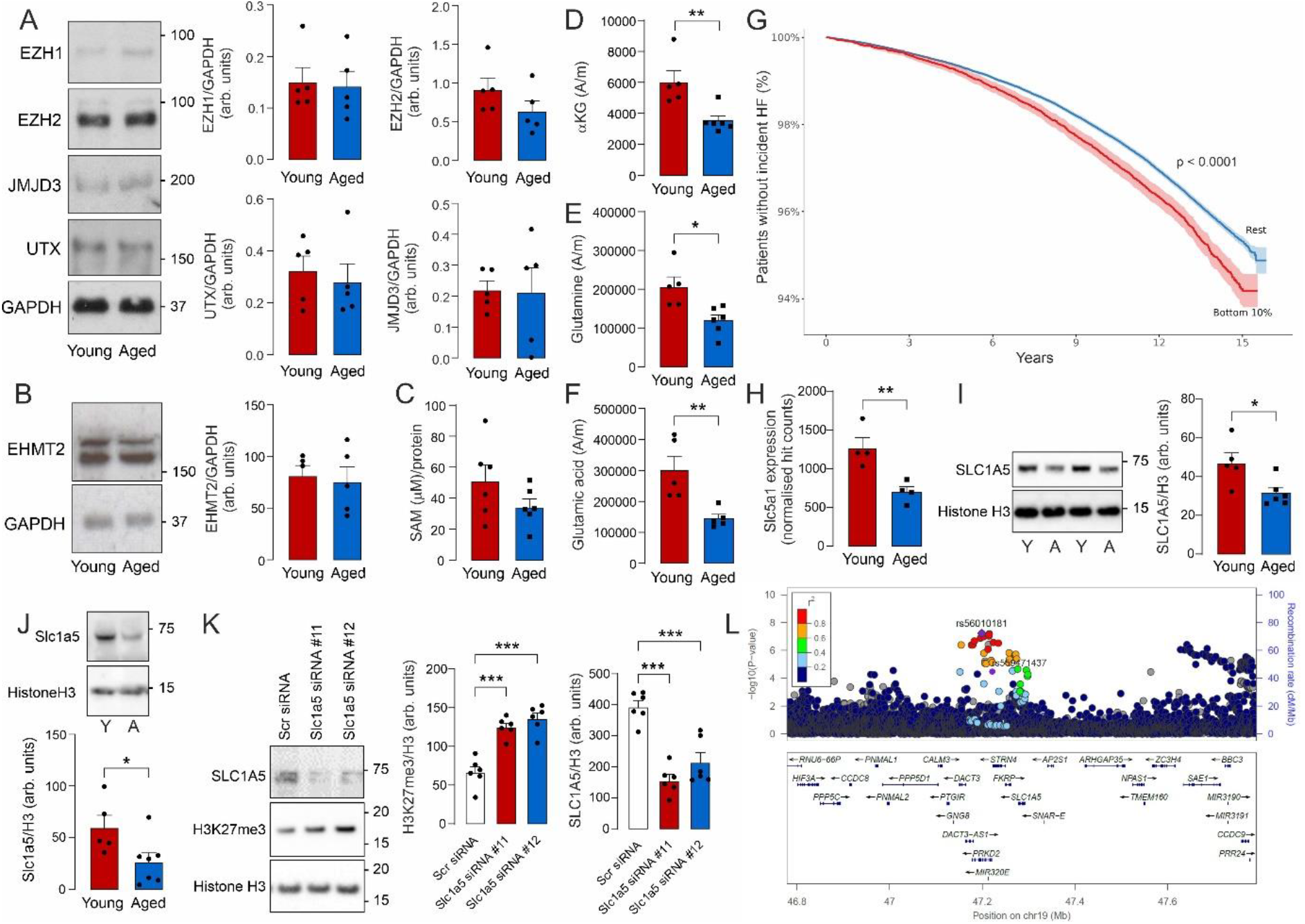
Increased H3K27me3 in aged mice is likely due to loss in glutamine transport. Expression of EZH1, EZH2, UTX and JMJD3 (A) or EHMT2 (B) in young and aged mice detected by immunblotting (n=5). C) The abundance of cardiac SAM in the young and aged mouse myocardium (n=6). The levels of alpha-ketoglutarate (D), glutamine (E) and glutamic acid (F) in the young and aged mouse myocardium (n=5). (G) Incidence of heart failure in patients with low (bottom 10%) vs higher circulating levels of glutamine. The expression of SLC1A5 in the young and aged mouse myocardium detected at the RNA (n=4) (H) and protein level (n=5) (I). J) The expression of SLC1A5 in aged human heart tissue detected by immunoblotting (n=5, 7). K) H3K27me3 in cardiomyocytes with or without knock-down of SLC1A5 (n=6). L) Locus Zoom Plot of the GWAS results of genetic variants within the scl5a1 gene region (±500kb) for their association with heart disease^22^. *P<0.05; **P<0.01; ***P<0.005 by Student *t* test or 1-way ANOVA with Tukey post hoc test (K).

### Elevation in H3K27me3 leads to metabolic reprogramming during aging

To further define the impact of elevated H3K27me3 on cardiac aging, genes harboring this modification were identified using CUT&RUN-seq and RNA-seq. Metabolism was identified as a major process likely to be affected in the aged myocardium due to the elevation in H3K27me3 (Fig. 3A, B). To substantiate the impact of H3K27me3 on metabolism, we assessed the effect of UTX knockdown on metabolite utilization in cardiomyocytes. Maximal respiration was found to be significantly reduced following UTX knockdown (Fig. 3C-E). Additionally, UTX knockdown led to a decrease in fatty acid utilization (Fig. 3E). Together, these findings suggest that the elevation of cardiac H3K27me3 during aging contributes to age associated changes in metabolism^24^.

**Figure 3.**
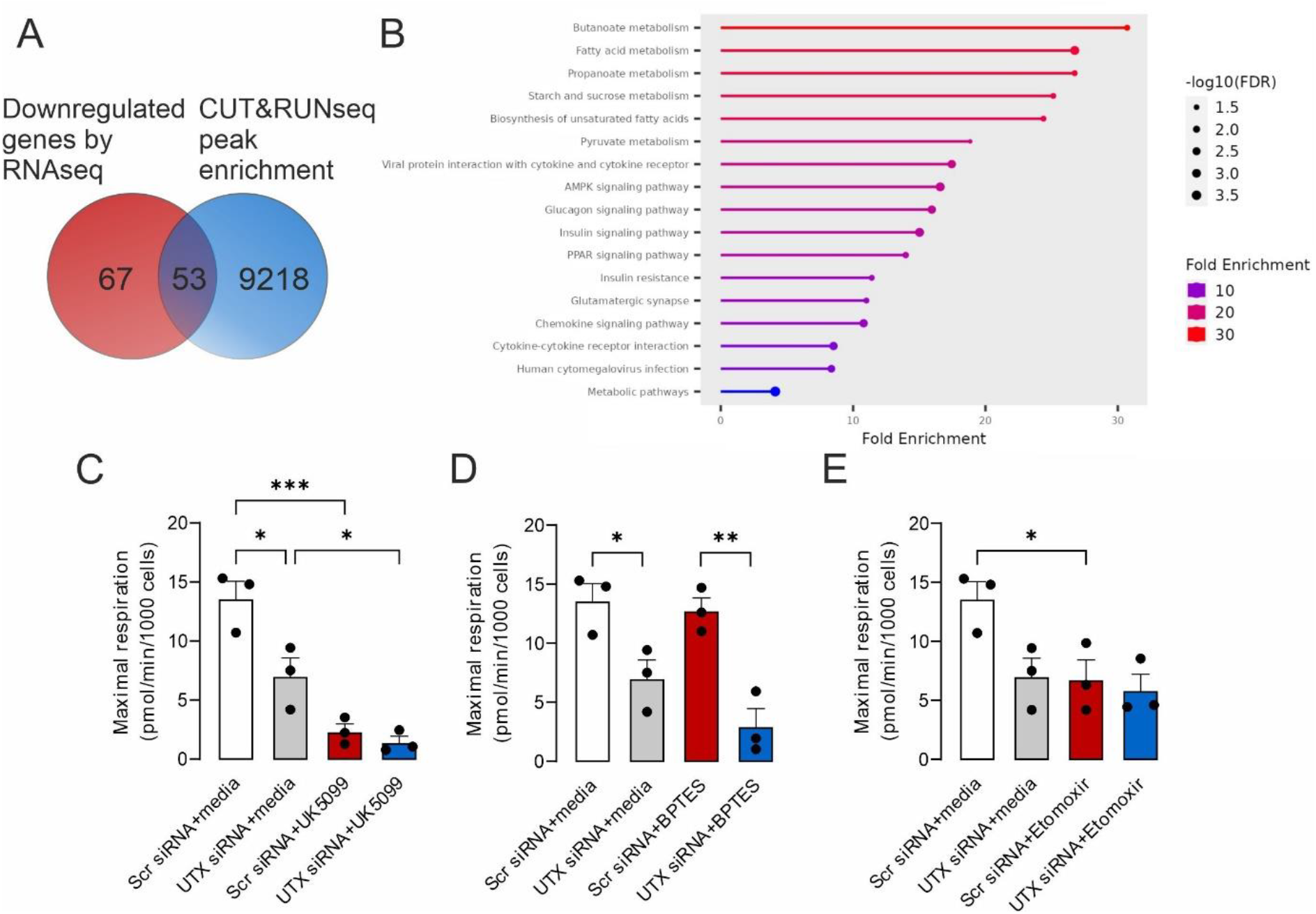
Elevation in H3K27me3 is likely to contribute to metabolic dysregulation during aging. A) Venn diagram of downregulated genes (RNA-seq) and CUT&RUN-seq enrichment for H3K27me3. B) Pathway analysis of common genes between RNA-seq and CUT&RUN-seq. Assessing the impact of UTX knock-down in cardiomyocytes on C) glycolysis, D) glutamine metabolism and E) fatty acid oxidation (n=3). *P<0.05; **P<0.01; ***P<0.005 by 1-way ANOVA with Tukey post hoc test.

### Dietary glutamine supplementation lowers H3K27me3 and improves function in the aged myocardium

To simulate the aging metabolic phenotype, glutamine abundance was restricted in cardiomyocyte culture media. This glutamine deprivation resulted in a significant increase in H3K27me3 levels in cardiomyocytes (Fig. 4A). Based on these findings, aged mice were fed a modified high-glutamine diet. This high-glutamine diet attenuated the increase in histone H3K27me3, an effect not observed in control or high-amino acid-fed mice (Fig. 4B). Furthermore, the high-glutamine diet led to significant improvements in cardiac function and reversed diastolic dysfunction in aged mice (Fig. 4C-H, Fig. S2). Notably, this effect was glutamine-dependent, as no similar improvements were seen in mice fed a high-amino acid diet. Consistent with the modified diet, glutamine and glutamic acid levels were significantly elevated in the hearts of aged mice fed a high-glutamine diet (Fig. 4I, J), with a trend toward increased α-KG (Fig. 4K).

**Figure 4.**
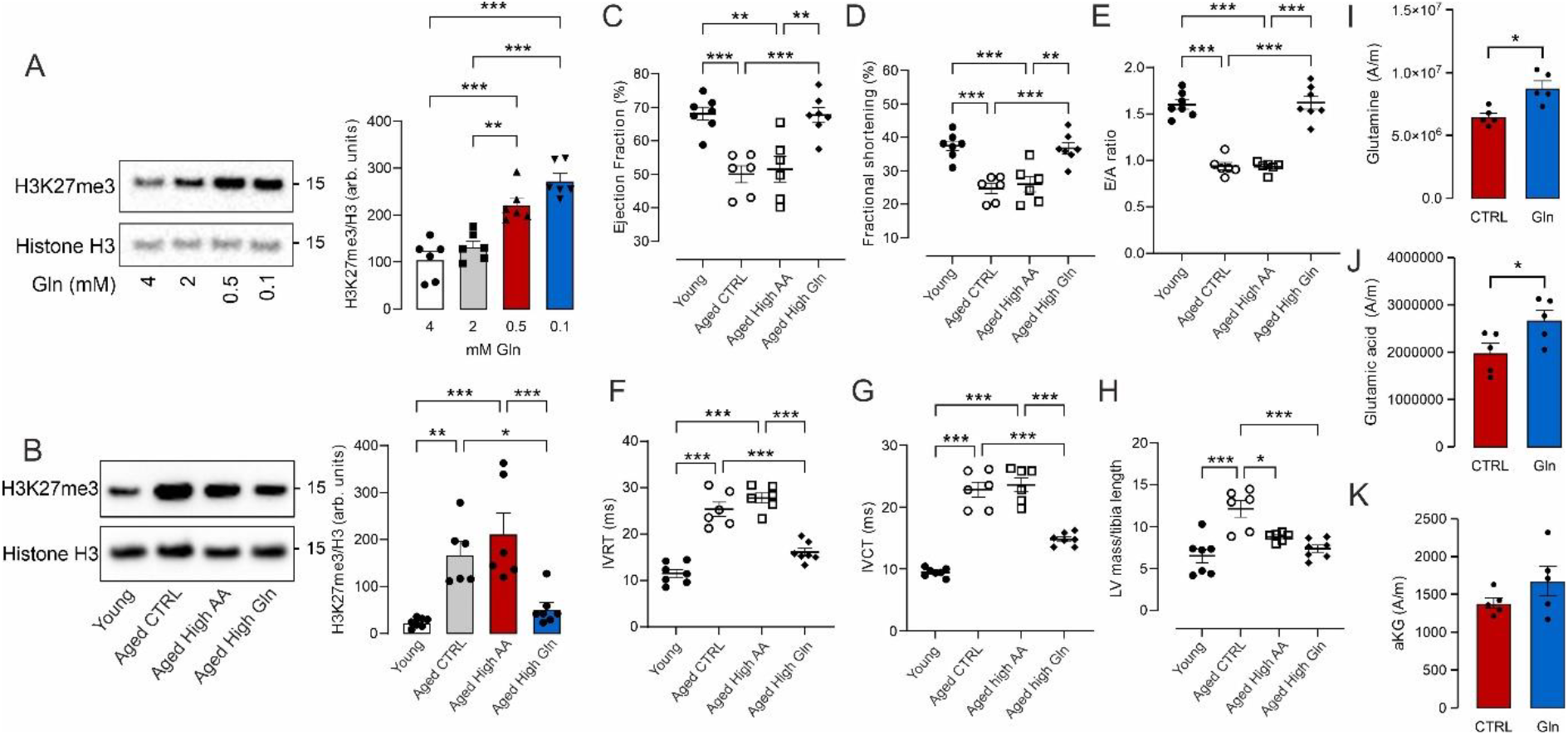
High glutamine diet lowers H3K27me3 and improves cardiac function. A) Measuring H3K27me3 in cardiomyocytes maintained in media with varying concentrations of glutamine (n=6). B) Myocardial H3K27me3 in young mice and aged mice fed a control (CTRL), high amino acid (AA) or high glutamine (Gln) diet (n=6, 7). Impact of age and diet on C) ejection fraction, D) fractional shortening, E) E/A, F) IVRT, G) IVCT and H) heart size in aged mice (n=6, 7). The levels of glutamine (I), glutamic acid (J), and alpha-ketoglutarate (K) in aged control and aged high glutamine fed mice measured used metabolomics (n=4) *P<0.05; **P<0.01; ***P<0.005 by 1-way ANOVA with Tukey post hoc test.

### Elevation in H3K27me3 leads to impairment in myocardial autophagy

Autophagy is a key biological process that protects the myocardium from aging. Since this catabolic process can be regulated by histone H3K27me3^14-18^, we further investigated this potential association. First, we established a cardiomyocyte model of elevated H3K27me3 by selectively knocking down the site-specific demethylase UTX. Loss of UTX resulted in a marked increase in H3K27me3 and a concurrent reduction in cardiomyocyte autophagy (Fig. 5A). This suggests that the increase in H3K27me3 during cardiac aging likely causes a loss of autophagy, contributing to impaired cellular and tissue function^25,26^. This hypothesis was supported by evidence of impaired autophagosome biogenesis during cardiac aging, which was improved in aged mice fed a high-glutamine diet (Fig. 5B). Consistent with enhanced autophagy and the removal of damaged cellular components, lipofuscin levels were lower in the myocardium of aged mice fed the high-glutamine diet (Fig. S3). These findings suggest that elevated cardiac H3K27me3 contributes to a decline in the cytoprotective process of autophagy during cardiac aging.

**Figure 5.**
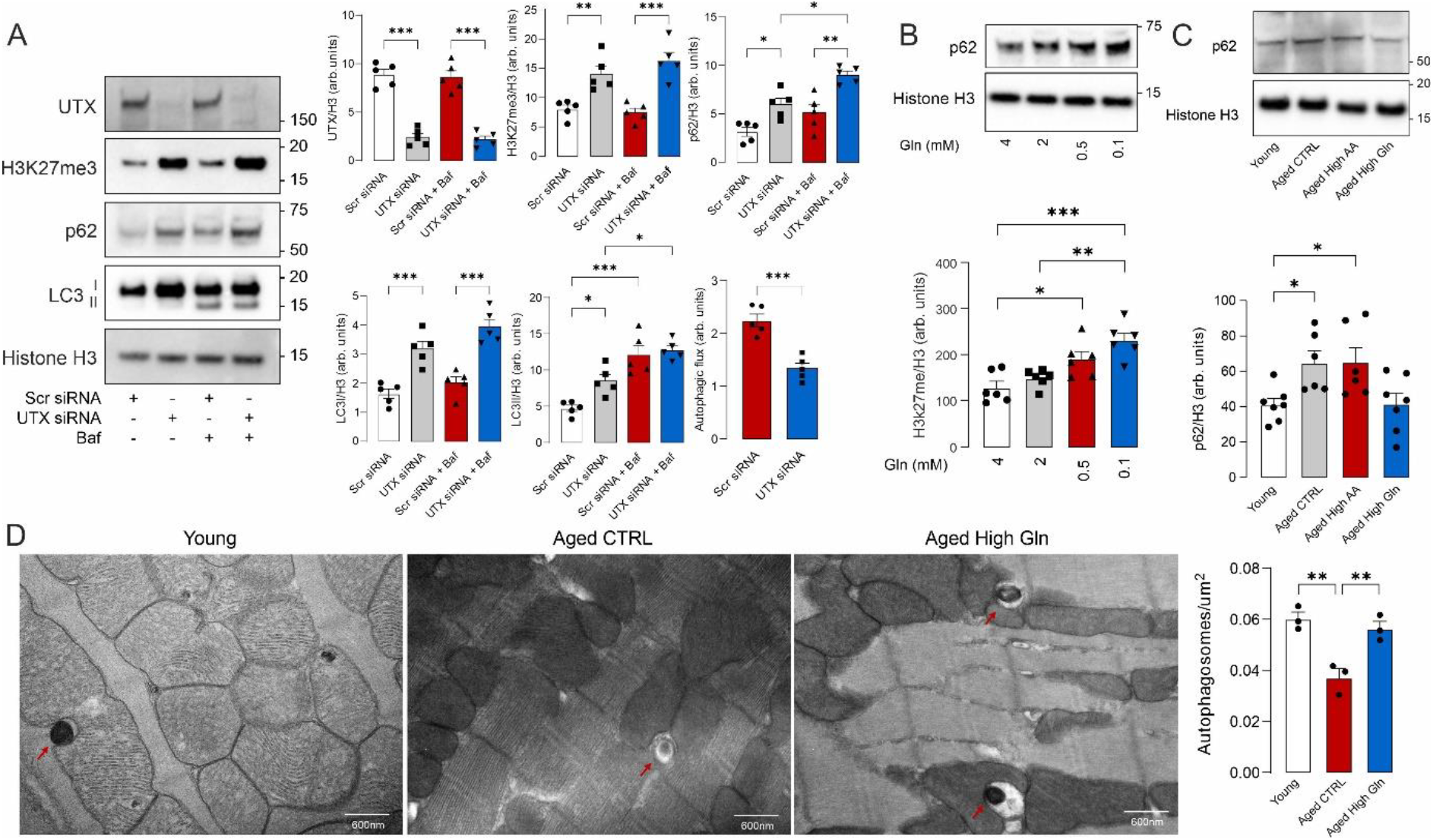
Elevation in H3K27me3 is likely to contribute to impaired autophagy during myocardial aging. A) H3K27me3, p62 and LC3 lipidation in cardiomyocytes with or without knock-down of UTX and detected by immunoblotting (n=5). B) Detection of p62 in cardiomyocytes maintained in media with different concentrations of glutamine (n=6). C) Detection of p62 in the myocardium of young mice and aged mice maintained on a control (CTRL), high amino acid (AA) or high glutamine (Gln) diet (n=6, 7). D) Analysis of autophagosome like structures in cardiac tissue from young and aged mice maintained on a control of high glutamine diet observed by electron microscopy (n=3). *P<0.05; **P<0.01; ***P<0.005 by 1-way ANOVA with Tukey post hoc test or Dunnett’s Test (C).

## Discussion

Our findings identify histone H3K27me3 as an epigenetic mark likely contributing to myocardial aging. Consistent with these findings, global changes in H3K27me3 have been previously observed in the heart during aging^27,28^. Additionally, this modification has been linked to health and lifespan. Heterozygous mutations in components of PRC2, specifically E(z) and Esc, decrease the global levels of histone H3K27me3 and extend lifespan in *Drosophila*^5,9^. This provides evidence that altering global levels of histone H3K27me3 can influence health. However, the complexity of this mark is underscored by its tissue-specific variability. Notably, the elevation in histone H3K27me3 during aging was restricted to the heart and brain. This finding aligns with observations in the brains of the senescence-accelerated mouse prone 8 (SAMP8) strain, a naturally occurring model of accelerated aging, where histone H3K27me3 was similarly increased^29^. The observation that histone H3K27me3 is specifically elevated in the aged heart and brain may be explained by the lack of cell division, leading to the accumulation of H3K27me3. This is consistent with the further increase of this mark observed in much older mouse myocardium. However, changes in metabolism and substrate availability may also contribute to the elevation of histone H3K27me3, as suggested by our findings that loss of glutamine metabolism likely mediates the increase in this epigenetic mark.

During aging, there was a loss of glutamine and α-ketoglutarate (αKG) in myocardial tissue. This depletion in glutamine metabolism is likely responsible for the increase in H3K27me3, as the demethylases UTX and JMJD3 require αKG as a cofactor^30^. This cofactor plays a crucial role in cardiovascular health and lifespan, likely mediated in part by its regulation of H3K27me3. Supplementation of αKG in mice has been shown to ameliorate age-related osteoporosis by regulating histone methylation, including H3K27me3^31^. Moreover, αKG supplementation also decreased frailty and extended lifespan in mice, with a significant improvement in cardiac function^32^.

The loss of glutamine metabolism during aging is likely attributed to a marked reduction in the expression of its transmembrane transporter, Slc1a5, in the aged myocardium. This finding aligns with the likely importance of Slc1a5 in maintaining cardiac health as single nucleotide polymorphisms within this gene showed evidence of association with heart disease. Furthermore, decrease in Slc1a5 expression is observed in heart failure patients, which results in impaired glutamine homeostasis^33^. Additionally, circulating glutamine levels was found to be inversely associated with the incidence of heart failure. This is consistent with previous studies where higher dietary glutamine intake was associated with a lower risk of mortality, particularly from cardiovascular causes, in both men and women^34^. Together with our findings this suggest that the decline in Slc1a5 expression and disrupted glutamine metabolism may be common features of cardiac dysfunction. This hypothesis is further supported by our feeding study, where glutamine supplementation preserved cardiac function and limited hypertrophy during aging.

The benefits of glutamine supplementation are likely mediated by its ability to attenuate the age-dependent elevation in H3K27me3. By limiting H3K27me3, glutamine supplementation is likely to improve the cytoprotective process of autophagy, which plays a key role in health and longevity^12^. Autophagy maintains cellular homeostasis by removing redundant and dysfunctional biomolecules and organelles, as well as providing bioenergetic intermediates during hypoxia and nutrient deprivation. In aged mice, evidence of impaired autophagy was observed, which was subsequently improved with dietary glutamine supplementation. Additionally, in cardiomyocytes, elevated H3K27me3 was associated with a loss of autophagic flux. This is consistent with previous findings linking histone H3K27me3 to the regulation of autophagy. For instance, changes in the distribution of H3K27me3 have been observed along autophagy-related genes such as LC3B, ATG4B, and p62/SQSTM1 during starvation^14^. Furthermore, chidamide, a benzamide inhibitor of histone deacetylases with potent antimyeloma activity, was shown to suppress autophagy by upregulating H4K16ac and H3K27me3 in the promoter regions of the autophagy-related gene LC3B^15^. Increased methyltransferase activity is also associated with suppressed autophagy. The methyltransferase EHMT2, for example, represses the expression of autophagy-associated proteins such as LC3B, WIPI1, and DOR, thus limiting this catabolic process^16^. Inhibition of EHMT2 by BIX01294 induces autophagy in human glioma cells by upregulating the expression of autophagy-related genes^17^. Moreover, the methyltransferase EZH2 epigenetically represses several negative regulators of the mTOR pathway, including TSC2, RHOA, DEPTOR, FKBP11, RGS16, and GPI^18^. The downregulation of TSC2 by EZH2 leads to mTOR activation, which subsequently inhibits autophagy. In our study when analysing processes likely to be regulated by H3K27me3 the AMPK pathway was detected. As AMPK plays a crucial role in regulating autophagy it is likely that H3K27me3-dependent suppression of genes within this pathway contribute to impairment in this catabolic process.

In addition to regulating autophagy, histone H3K27me3 is likely to impact myocardial aging by altering cellular metabolism. This is consistent with the identification of genes involved in metabolic processes with expression that was likely impacted by histone H3K27me3 during aging. Furthermore, elevating H3K27me3 in cardiomyocytes resulted in a reduction in maximal respiration and a loss in fatty acid utilisation. These findings align with the aged mouse and human myocardium, where there is limited metabolic flexibility and an increased shift toward glycolysis rather than fatty acid oxidation^12^. Consistent with our results, histone H3K27me3 is known to impact metabolic processes^35^, including the modulation of glycolysis during aging in *Drosophila*^5^.

Together, our findings identify myocardial H3K27me3 as a key process altered during aging, with its elevation likely contributing to impaired autophagy and metabolic reprogramming. Targeting this process, including through dietary supplementation of glutamine, holds potential for preserving cardiac function in the aging population.

## Sources of Funding

This study was supported by a Mark Braimbridge Cardiovascular PhD Studentship, King’s BHF Centre of Research Excellence (RE/18/2/34213; RE/24/130035), British Heart Foundation (PG/22/10932, PG/24/11731, IA/F/23/275048) and the Medical Research Council (MR/Y01185/1).

## Disclosures

None.

## Notes

### Competing Interest Statement

The authors have declared no competing interest.

